# Elimination of PCR duplicates in RNA-seq and small RNA-seq using unique molecular identifiers

**DOI:** 10.1101/251892

**Authors:** Yu Fu, Pei-Hsuan Wu, Timothy Beane, Phillip D. Zamore, Zhiping Weng

**Author notes:** These authors contribute equally to this work (Yu Fu, Pei-Hsuan Wu). **For correspondence:** (ZW), (PDZ).

## Abstract

RNA-seq and small RNA-seq are powerful, quantitative tools to study gene regulation and function. Common high-throughput sequencing methods rely on polymerase chain reaction (PCR) to expand the starting material, but not every molecule amplifies equally, causing some to be overrepresented. Unique molecular identifiers (UMIs) can be used to distinguish undesirable PCR duplicates derived from a single molecule and identical but biologically meaningful reads from different molecules. We have incorporated UMIs into RNA-seq and small RNA-seq protocols and developed tools to analyze the resulting data. Our UMIs contain stretches of random nucleotides whose lengths sufficiently capture diverse molecule species in both RNA-seq and small RNA-seq libraries generated from mouse testis. Our approach yields high-quality data while allowing unique tagging of all molecules in high-depth libraries. Using simulated and real datasets, we demonstrate that our methods increase the reproducibility of RNA-seq and small RNA-seq data. Notably, we find that the amount of starting material and sequencing depth, but not the number of PCR cycles, determine PCR duplicate frequency. Finally, we show that computational removal of PCR duplicates based only on their mapping coordinates introduces substantial bias into data analysis.

## Introduction

High-throughput sequencing of long (>100 nt) or small (18–50 nt) RNA provides a quantitative measure of RNA abundance. However, RNA-seq and small RNA-seq library construction can introduce bias at multiple steps, such as fragmentation of long RNAs, reverse transcription, adapter ligation, library amplification by PCR, and sequencing. Commonly used high-throughput sequencing platforms, including those made by Illumina and Pacific Biosciences, require PCR amplification during library construction to increase the number of cDNA molecules to an amount sufficient for sequencing. However, PCR stochastically introduces errors that can propagate to later cycles (Cha and Thilly, 1993; Dohm et al., 2008). PCR also amplifies different molecules with unequal probabilities (Cha and Thilly, 1993). PCR duplicates are reads that are made from the same original cDNA molecule via PCR.

A common practice to eliminate PCR duplicates is to remove all but one read of identical sequences, assuming that such reads have been created from the same cDNA molecule by PCR (Li et al., 2009). This assumption may be flawed, especially with ever higher sequencing throughput, which increases the chance of observing reads with identical sequences but from different cDNA molecules. The situation is further exacerbated for small genomes and for techniques that interrogate a subspace of the genome. For example, the majority of small RNA-seq reads are microRNAs (miRNAs) or PIWI-interacting RNAs (piRNAs), which derive from loci that amount to just a few percent of the genome (Aravin et al., 2006; Girard et al., 2006; Brennecke et al., 2007; Li et al., 2013). The assumption also has systematic biases. For example, a shorter gene is more likely to give rise to identical RNA-seq reads than a longer gene with the same transcript level, simply because the “genomic space” for the random process of RNA fragmentation is smaller for the shorter gene. Finally, the conventional definition of PCR duplicates is based on mapping coordinates—reads mapping to the exact same genomic location are considered to have identical sequences. However, many small RNAs with the same sequence can be produced from multiple genomic loci; thus, strategies using genome mapping to identify PCR duplicates ignore the situation that identical reads arise from distinct sites in the genome.

Standard library preparation and sequencing procedures typically have prespecified PCR and sequencing error rates, but parameters such as the amount of starting RNA used to generate a library, the number of reads sequenced (i.e., sequencing depth), and the number of PCR cycles used are often adjusted to accommodate sample source, abundance, and quality. While the notion that more PCR amplification increases artefactual duplicate reads in high-throughput sequencing makes intuitive sense and is widely accepted, high PCR cycle numbers are often necessitated by scarce starting materials, another likely cause for duplicate reads. Thus, the contribution of PCR cycle number to PCR duplicates is often confounded with the contributions of starting materials and sequence depth.

Unique molecular identifiers (UMIs) are often used to accurately detect PCR duplicates accurately and quantify transcript abundance (Fu et al., 2011; Kivioja et al., 2011; Shiroguchi et al., 2012; Fu et al., 2014a; Fu et al., 2014b; Islam et al., 2014; Collins et al., 2015; Smith et al., 2017). If each molecule in the starting pool is barcoded with a UMI, i.e., all molecules are unique, then reads with the same UMI must be PCR duplicates. In practice, only the molecules in the starting pool that have identical sequences need to have different UMIs.

One strategy to incorporate UMIs introduces pre-defined, manually-selected sequences into the adapters. This strategy can avoid UMIs with suboptimal GC content and minimize complementarity between or within UMI sequences (Shiroguchi et al., 2012). Because UMI identities are unambiguously defined, sequencing and PCR errors can be easily corrected. However, implementing pre-defined UMIs requires a large number of costly, custom-synthesized oligonucleotides.

An alternative strategy employs adapters that contain random nucleotides at certain positions in the adapters. The combinations of the random-nucleotide positions lead to an exponential number of different UMIs at almost no extra cost, because incorporating a random nucleotide costs the same as incorporating a specific nucleotide during DNA synthesis. UMIs bearing either five (4^5^ = 1,024 unique barcodes) or ten random nucleotides (4^10^ = 1,048,576 unique barcodes) were implemented cost-effectively and shown to improve PCR duplicate removal (Kivioja et al., 2011; Islam et al., 2014). A higher number of unique combinations can be achieved simply by increasing the number of random-nucleotide positions. The number of UMI combinations must be sufficiently large because as mentioned above, the chance that two cDNA molecules with identical sequences in the starting pool are tagged with the same UMI combination needs to be infinitesimally small.

Here, we describe novel experimental protocols and computational methods to unambiguously identify PCR duplicates in RNA-seq and small RNA-seq data. We show that removing PCR duplicates using UMI information is accurate, whereas removing PCR duplicates without UMIs is overly aggressive, eliminating many biologically meaningful reads. Finally, we show that the amount of starting materials and sequencing depth determine the level of PCR duplicates, without additional contribution from the extent of PCR amplification.

## Results

### Adapting standard RNA-seq procedures to incorporate UMIs

To incorporate UMIs into RNA-seq, we modified a published, strand-specific, library construction protocol (Zhang et al., 2012). The original method has proved to be robust and time-efficient, and the adapter ligation step uses DNA adapter oligonucleotides that can be readily synthesized at a low cost (Li et al., 2013; Hayashi et al., 2014; Mohn et al., 2014; Zhang et al., 2014). The standard protocol uses a single Y-shaped DNA adapter comprising two partially complementary oligonucleotides and an unpaired 3′ thymidine that pairs with the single adenine tail added to both ends of the double-stranded cDNA fragments. We modified the adapters by inserting a five-nucleotide random UMI (Fig. 1A,B). Consequently, each cDNA fragment is ligated to an adapter with a UMI at each end, randomly choosing one out of 1,048,576 (4^5^ × 4^5^) possible combinations provided by two UMIs.

**Figure 1.**
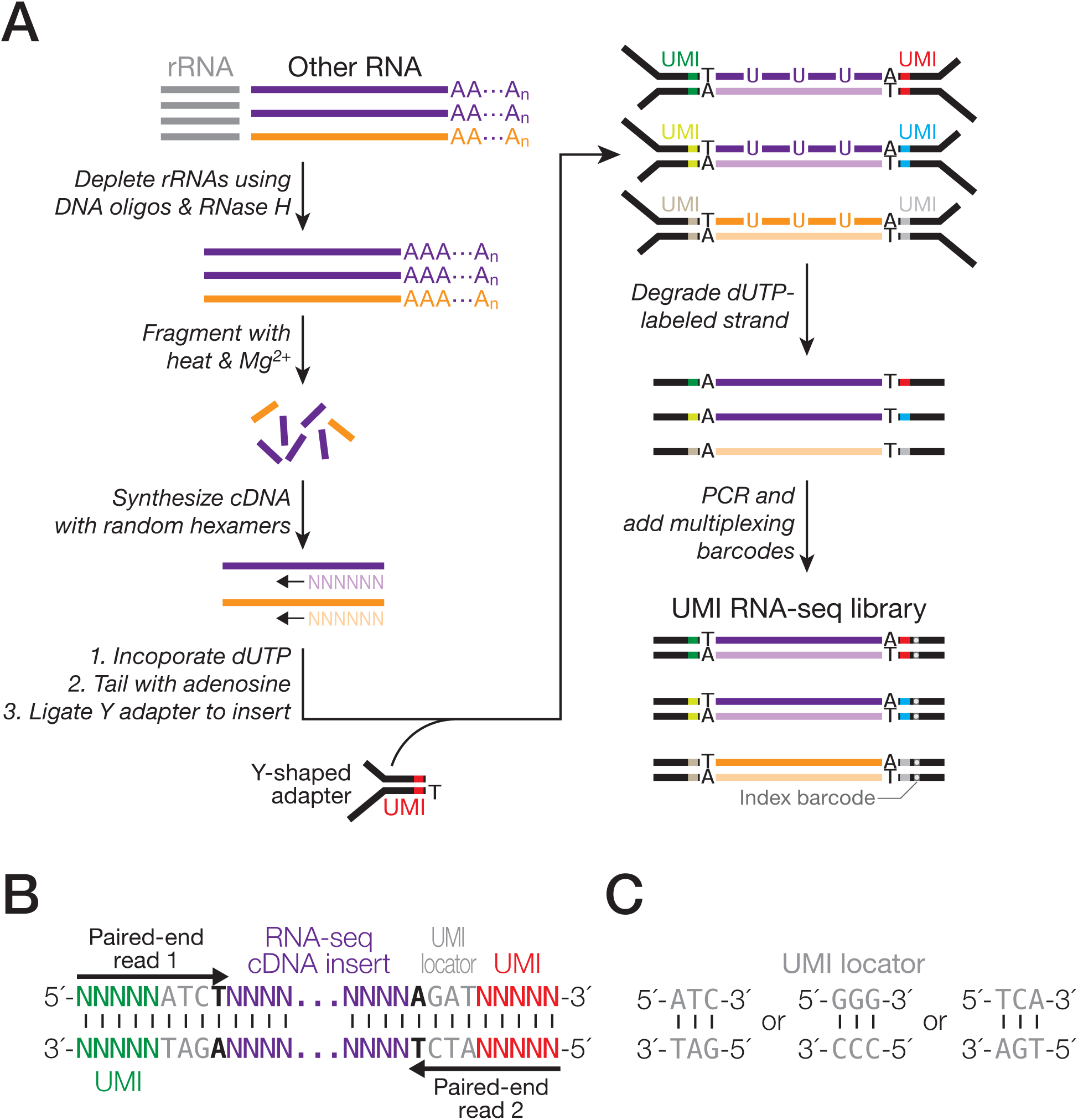
UMI incorporation into RNA-seq. (A) Overall workflow. Schematic of a read produced from RNA-seq with UMIs (B) and of UMI locators (C).

Our UMI RNA-seq adapters were designed so that the sequencing reaction begins at the very first nucleotide of the 5′ UMI (Fig. 1B). The random nucleotides of UMIs offer the sequence diversity in the initial five sequencing cycles. This sequence diversity is critical for commonly used Illumina sequencing platforms, such as HiSeq, MiSeq, and NextSeq, to generate base-calling templates and make accurate models for discriminating read clusters (Illumina, 2014; Mitra et al., 2015). To avoid insertions or deletions within or flanking a UMI, albeit rare, from altering the UMI identity, we further designed a “UMI locator”, a pre-defined trinucleotide 3′ to the UMI (e.g. 5′– NNNNNATC–3′). The trinucleotide serves as an anchor allowing unambiguous identification of each UMI (Fig. 1B). Taking the properties of our sequencing instrument of choice—NextSeq 500—into consideration, the 3 nt UMI locator sequence and the mandatory thymidine required for ligation that immediately follows (Fig. 1B) corresponded to the sequencing cycles 6–9, after the first five critical cycles required by the instrument for template generation (Illumina, 2016). However, NextSeq still deemed these four invariant positions of low complexity and reported low-quality sequencing data. Previous approaches to tackle this problem include increasing the diversity of the initial sequences in the library, mixing the library with a high diversity sample (spike-in), lowering sequencing cluster density, or any combination of the above (Mitra et al., 2015). We designed three UMI locator sequences (Fig. 1C), and, by pooling adapters with one of these sequences at equimolar amounts, we were able to resolve the low complexity problem. Using this approach, we generated RNA-seq libraries from mouse brain, heart, kidney, liver, lung, muscle, spleen, and testis total RNAs. The libraries were sequenced at a read depth, coverage, and quality comparable to libraries generated using the original protocol without UMIs (Supplemental Table S1). Thus, our method of incorporating UMIs, as well as UMI locator, does not interfere with library preparation and sequencing. We subsequently observed that even two different UMI locator sequences sufficed to overcome the erroneous low-quality calling by NextSeq (small RNA-seq, Fig. 2).

**Figure 2.**
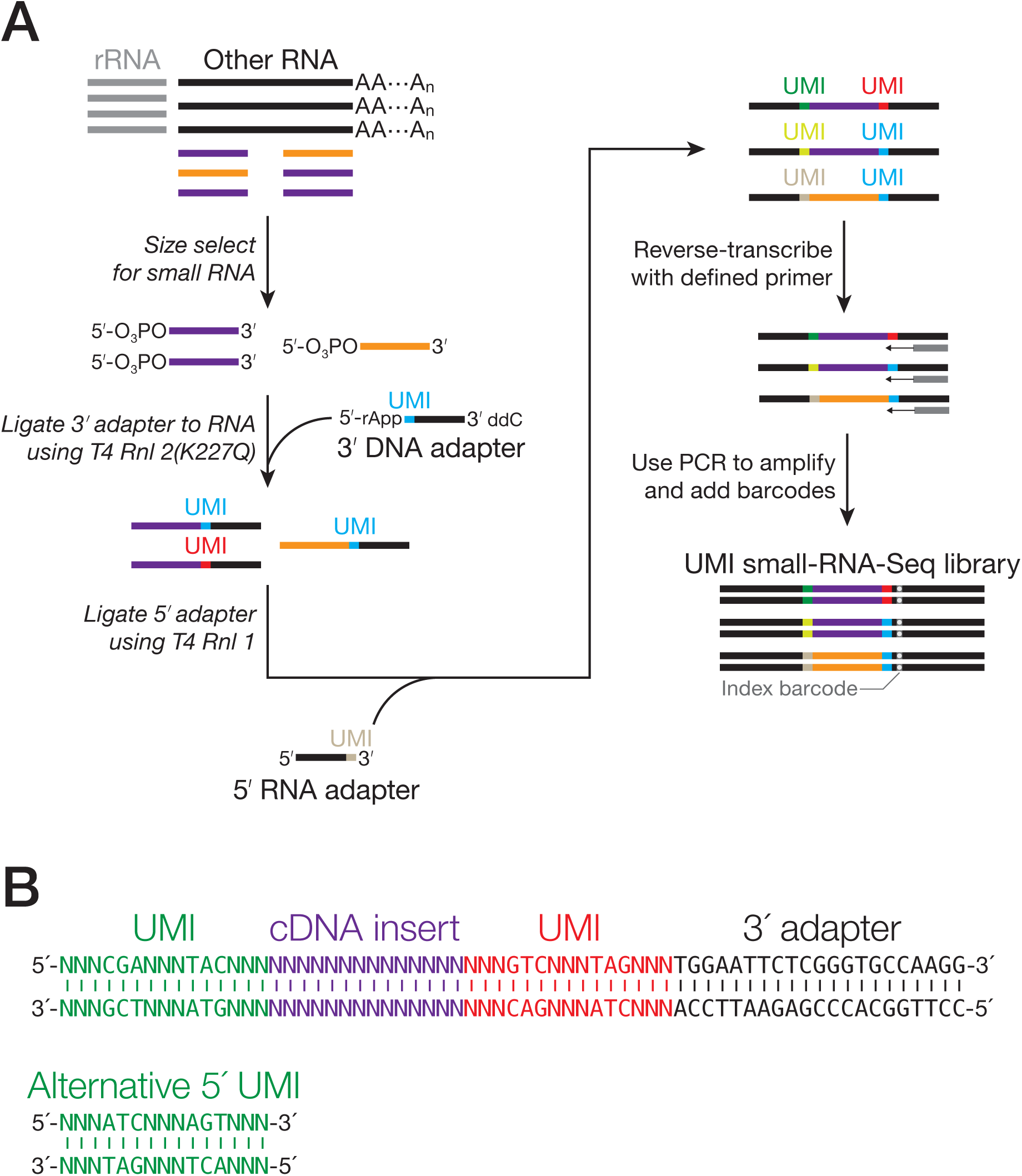
UMI incorporation into small RNA-seq. (A) Overall workflow. The method uses a 3′ adapter composed of DNA, except for a single, 5′ ribonucleotide (rA); the 5′ adapter is entirely RNA. A standard index barcode allows multiplexing. (B) Schematic of a read produced from small RNA-seq with UMIs.

### Adapting standard small RNA-seq protocol to incorporate UMIs

Previously, we established a reliable and robust small RNA-seq protocol by modifying a published method which utilizes oligonucleotides compatible with Illumina sequencing platforms (Lau et al., 2001). Compared to UMI RNA-seq, incorporation of UMIs into this small RNA-seq protocol requires additional considerations. First, the number of distinct UMI combinations needs to be significantly greater than what is required for RNA-seq. For example, millions of piRNA species—an abundant class of small RNAs in the animal germ line—can be routinely detected in a single individual, and it is estimated that there can be as many as 1 million distinct piRNA molecules in a single spermatocyte or round spermatid (Girard et al., 2006; Aravin et al., 2007; Brennecke et al., 2007; Houwing et al., 2007). The most abundant piRNA species in this study has 42,281 reads. In the soma, the most abundant miRNA can take up >40% of the total sequencing depth (Yue et al., 2014) —tens of millions of reads. Such enormous abundance requires a sufficiently high number of UMI combinations to capture all distinct sequences. Second, the length range of small RNAs (< 50 nt) plus a longer UMI is still well within the read length achievable by common sequencing instruments. Third, the length of a small RNA is a defining feature of its identity and thus, insertions or deletions could lead to misclassification of small RNAs. The second and third considerations also indicate that small RNA-seq is ideally suited for the testing of a large combination of UMIs.

We tested UMIs containing 10 consecutive random nucleotides. Although both the 3′ and 5′ adapters containing 10-nt UMIs ligated to small RNAs with nearly the same efficiency as the original adapters without UMIs, the resulting small RNA-seq libraries yielded unexpectedly short, variable-length reads that contained truncated insert and adapter sequences (data not shown). We speculate that long stretches of random nucleotides interfere with oligonucleotide annealing, a critical step in cDNA synthesis, PCR, and sequencing, by increasing the chance that a primer anneals to a UMI instead of its target sequences. Inter‐ and intramolecular annealing of 10 nt UMIs may also contribute to truncated reads.

To avoid a long stretch of random nucleotides, we used the UMI locator strategy described above to space out several short stretches of random nucleotides. For each adapter, we designed three trinucleotide UMI sequences, each separated from another by a trinucleotide UMI locator (e.g., 5′–NNN-CGA-NNN-TAC-NNN–3′; Fig. 2A,B). Two adapters with such UMIs can produce a trillion combinations, which should suffice all deep-sequencing applications. Similar to our RNA-seq strategy, we designed adapters with two different sets of UMI locator sequences—mixed at equimolar—to increase the sequence complexity in the early sequencing cycles. This strategy allowed us to successfully generate and sequence the UMI small RNA-seq libraries, unambiguously locate UMIs, and computationally remove reads containing insertions or deletions in UMIs due to reverse transcription, PCR, and sequencing errors (Fig. 2B). We tested our method using total RNAs extracted from mouse testes isolated 17.5 days after birth. To assess the impact of the amount of starting materials on PCR duplicates, we prepared small RNA-seq libraries using a range of 39–5,000 ng RNAs made from serial dilution. To test the effect of PCR cycles, we gradually increased the PCR cycles for each library with a two-cycle increment. The resulting UMI small RNA-seq libraries yielded high-quality sequencing data, comparable to those generated with the original non-UMI protocol (Supplemental Table S1).

### Diverse UMIs capture all read species in RNA-seq and small RNA-seq

As mentioned above, to accurately identify PCR duplicates using UMIs, it is critical that the number of distinct UMIs far exceeds the maximal number of starting molecules with identical sequences, such that these molecules have an infinitesimal probability of being ligated to adapters with the same UMI. Previous UMI methods were designed for sequencing single cells or an organism with a less complex transcriptome than mammals (Shiroguchi et al., 2012; Fu et al., 2014a). In particular, testis has a higher-complexity transcriptome than many other tissues such as muscle, liver, and even brain (Soumillon et al., 2013), demanding a large number of UMI combinations. Our UMI RNA-seq protocol theoretically provides ∼1 million (4^10^) distinct combinations, and we asked whether this diversity far exceeded the maximal number of reads with identical sequences in our libraries. Indeed, the transcripts derived from the 299-bp *7S RNA 1* gene produce 19,271 identical reads mapping to the same genomic coordinate, all of which are attached to distinct UMI sequences, indicating that all of these reads were from different starting RNA molecules. In conclusion, our UMI RNA-seq protocol is more than sufficient to disambiguate biologically identical reads from PCR duplicates.

Our UMI small RNA-seq provides an even higher number of possible combinations with 18 nt UMIs—68.7 billion (4^18^)—much larger than the number of reads currently produced by a sequencing run. The most abundant small RNA species in our datasets is a piRNA with 42,281 reads, far fewer than the number of UMI combinations our protocol provides. We conclude that the UMI lengths used in the RNA-seq and small RNA-seq protocols contain a sufficient UMI diversity for current and, most likely, future sequencing experiments.

### Error-correction for UMIs only slightly improves PCR duplicate identification

To test whether UMIs could accurately identify PCR duplicates, we first evaluated their performance using simulated data. Assuming a library has sufficiently diverse UMI sequences, the simplest way to determine biologically identical reads is to look for reads with the same sequence but are tagged by different UMIs. This approach assumes that there is no error in the replication or reading of the UMI sequences, since such errors could render identical UMI sequences different and vice versa, causing misidentification of PCR duplicates. UMI errors could occur during PCR sequencing, and computationally correcting these errors has been shown to improve identification of PCR duplicates (Islam et al., 2014; Bose et al., 2015; Macosko et al., 2015; Yaari and Kleinstein, 2015; Smith et al., 2017).

We designed a strategy for correcting UMI errors with the following considerations in mind. First, UMI errors are rare, with rates stipulated by the chemistry of PCR and sequencing (∼10^-5^ and ∼10^-3^ errors per position respectively) (Lundberg et al., 1991; Zhou et al., 1991; Flaman et al., 1994; Schirmer et al., 2016)). Second, when two sufficiently long UMIs (for example, 10 and 18 nt in this study) that differ by just one base are connected to two reads with identical sequences, the probability that these are PCR duplicates of the same UMI with an error, albeit low (*p* < 10^-3^) is still much higher than the probability that these are two distinct UMIs (*p* = 4^-10^ for RNA-seq and 4^-18^ for small RNA-seq in this study). Adopting an error-correction method previously developed for RNA-seq (Smith et al., 2017), we built a UMI graph for each group of reads (Fig. 3A). For RNA-seq, the reads that map to the same genomic position form a group. This approach does not work for small RNAs, because they often originate from multiple genomic loci. Thus, we simply defined a group of small RNA reads as those with identical sequences. In both the RNA-seq and small RNA-seq UMI graphs, a node denotes a unique UMI and further holds the number of reads with that UMI (Fig. 3A). For each pair of UMIs (say, UMI *a* and UMI *b*) that differ by just one base (one edit distance apart), we connect their nodes if *n*_a_ ≥ 2 × *n*_b_ − 1, where *n*_a_ and *n*_b_ represent read counts for the two UMIs. We require a twofold difference between *n*_a_ and *n*_b_, because as described above, the error rates for PCR and sequencing are low, and the twofold differences corresponds to the most extreme case whereby an error occurred during the first PCR cycle. However, a twofold difference is too stringent for pairs of UMIs with low read counts (e.g., 1 versus 2), for which the error predominantly arose from sequencing. We therefore added “−1” to ensure that these UMIs could be connected. All connected UMIs are then assumed to originate from the most abundant UMIs in the graph. This scheme allows correction of two or more errors in UMIs, provided that the intermediate UMIs are observed (for example, the intermediate UMI with one error and UMI with two errors in Fig. 3A–B). One could relax the stringency of this method by adding direct connections between two nodes that differ in two or more positions.

**Figure 3.**
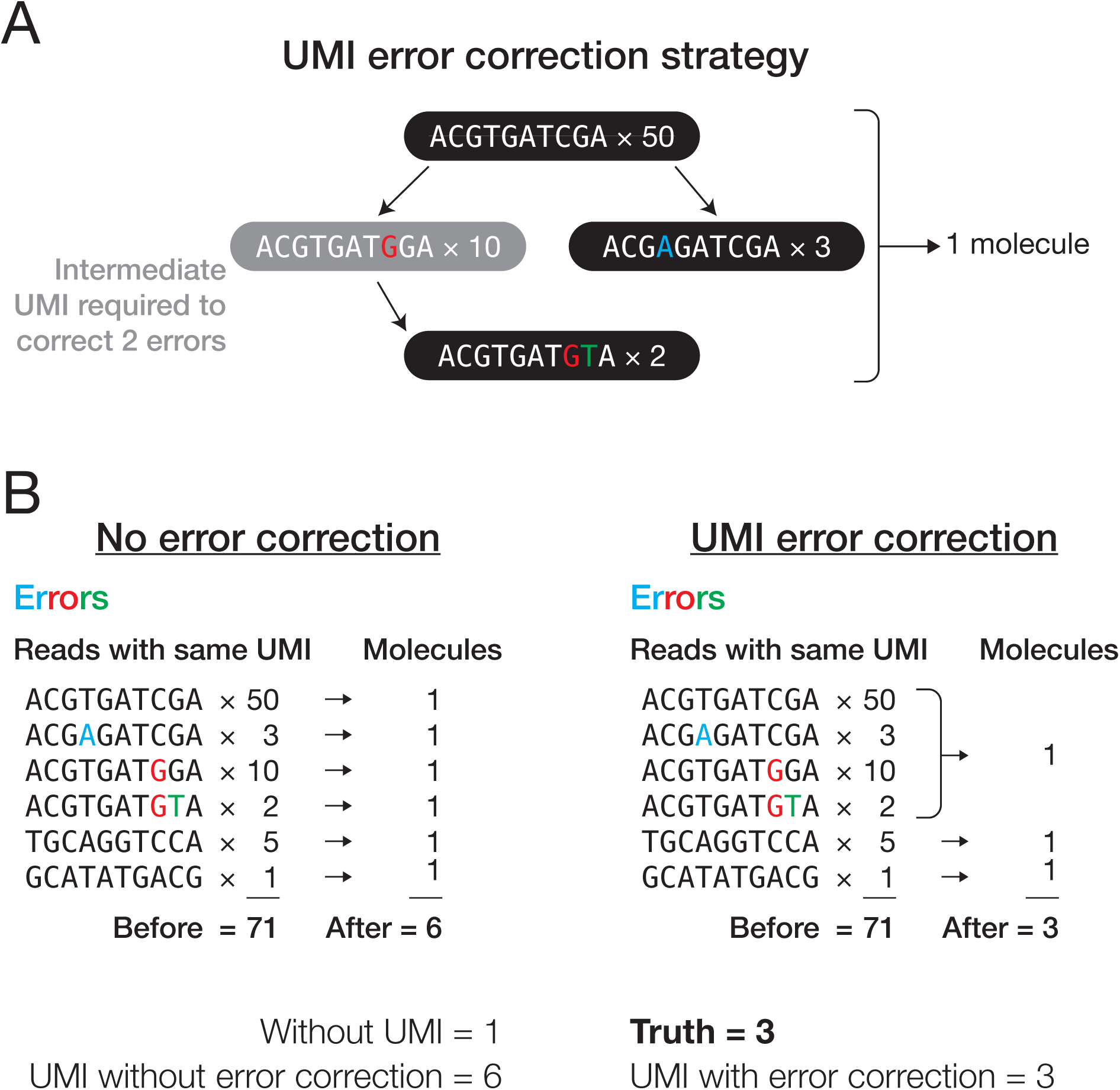
Identifying PCR duplicates. (A) Strategy for correcting errors in UMIs. (B) Illustration of how correcting errors in UMIs increases accuracy of PCR duplicate elimination.

The need for error-correction might depend on the experimental conditions, including the PCR amplification probability, PCR and sequencing error rates, UMI length, number of initial molecules, number of sequenced molecules, and number of PCR cycles. We performed computer simulations to investigate the effects of these seven experimental conditions on UMI error correction by systematically varying one variable at a time while holding the other six constant. Each round of simulation produced a known number of PCR duplicates and therefore, unlike experimental data, the true fraction of all reads corresponding to PCR duplicates can be determined in the simulated data. To assess the accuracy of PCR duplicate identification using UMIs, we calculated the difference between the number of reads after PCR duplicate removal (“estimate”) and the true value (“truth”) relative to the true value: (estimate − truth) / truth. This metric reflects the extent to which UMIs over- or underestimate the truth as a fraction of the true value. We started the simulation with 100 initial molecules. We then performed PCR by randomly assigning a probability to each molecule (tagged with an 18 nt UMI) to be duplicated in each PCR cycle. The probability follows a uniform distribution between *m* and 1, where *m* denotes minimum amplification probability (it can be any value between 0 and 1 and is set to 0.8 in the baseline condition). Minimum amplification probability can be interpreted as PCR efficiency, because the efficiency (average probability) that a molecule is doubled during each PCR cycle is (1-*m*)/2. Ten cycles of PCR (PCR error rate set to 3×10^-5^) (Lundberg et al., 1991; Zhou et al., 1991; Flaman et al., 1994) generated a pool of 61,000 ± 1,000 (mean ± S.D.) molecules. To test the effect of sequencing depth, we randomly drew 100 molecules from the pool for sequencing (sequencing error rate set to 10^-3^) (Schirmer et al., 2016) (Fig. 4; Supplemental Fig. S1). We call this set of parameters “baseline condition”, and it forms the base line from which we systematically varied each parameter. For each condition, we performed 10,000 trials.

**Figure 4.**
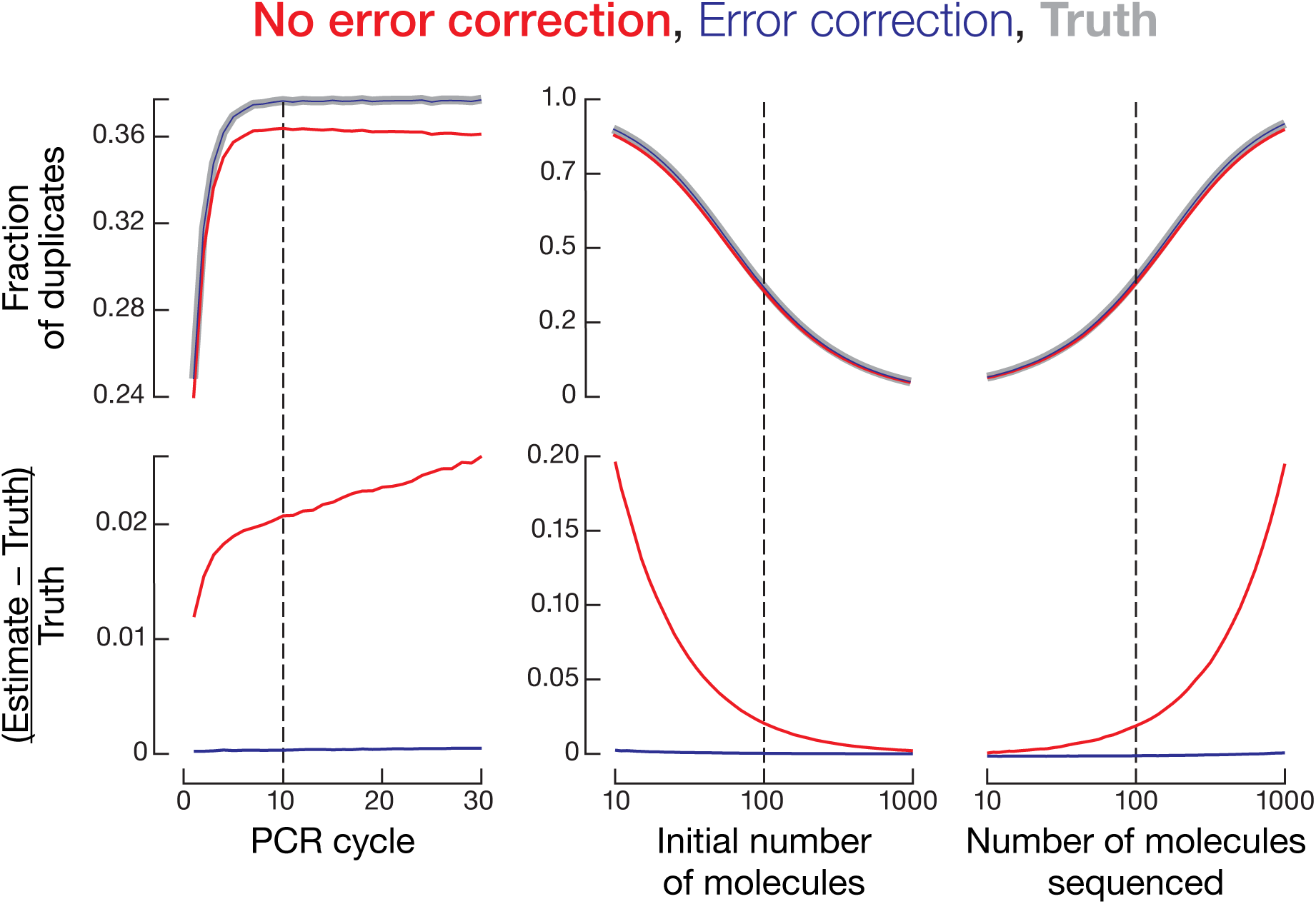
Simulation of PCR duplicate removal with or without error correction for UMIs. One parameter (PCR cycle number, starting material, or sequencing depth) was varied with the other parameters kept constant. Upper plots show the fraction of duplicates, while lower plots show the accuracy of duplicate detection. Each dotted line indicates the value for this parameter used in other simulations.

We first assumed that there was no error in UMIs (Fig. 3A) and found that on average, (estimate – truth) / truth = 2.10% across 10,000 trials under the baseline condition. Thus, without performing UMI error correction, we slightly overestimated the total number of biological molecules as an error in a UMI would artificially create an extra UMI, and in turn, we slightly underestimated the fraction of PCR duplicates (red vs gray lines in Fig. 4; Supplemental Fig. S1). Next, we used the UMI graph approach described above (Fig. 3A,B) for correcting errors in UMIs, and the new average of (estimate – truth) / truth = 0.0388%. Even though correcting UMI errors consistently gives better (estimate – truth) / truth than not correcting the errors, the absolute difference in the fractions of PCR duplicates between the two approaches is small (Fig. 4; Supplemental Fig. S1). For example, under the baseline condition, the true fraction of duplicates was 37.8 ± 3.2%; without correcting UMI errors yielded 36.5 ± 3.3%, and correcting UMI errors gave 37.8 ± 3.2%.

**Figure S1.**
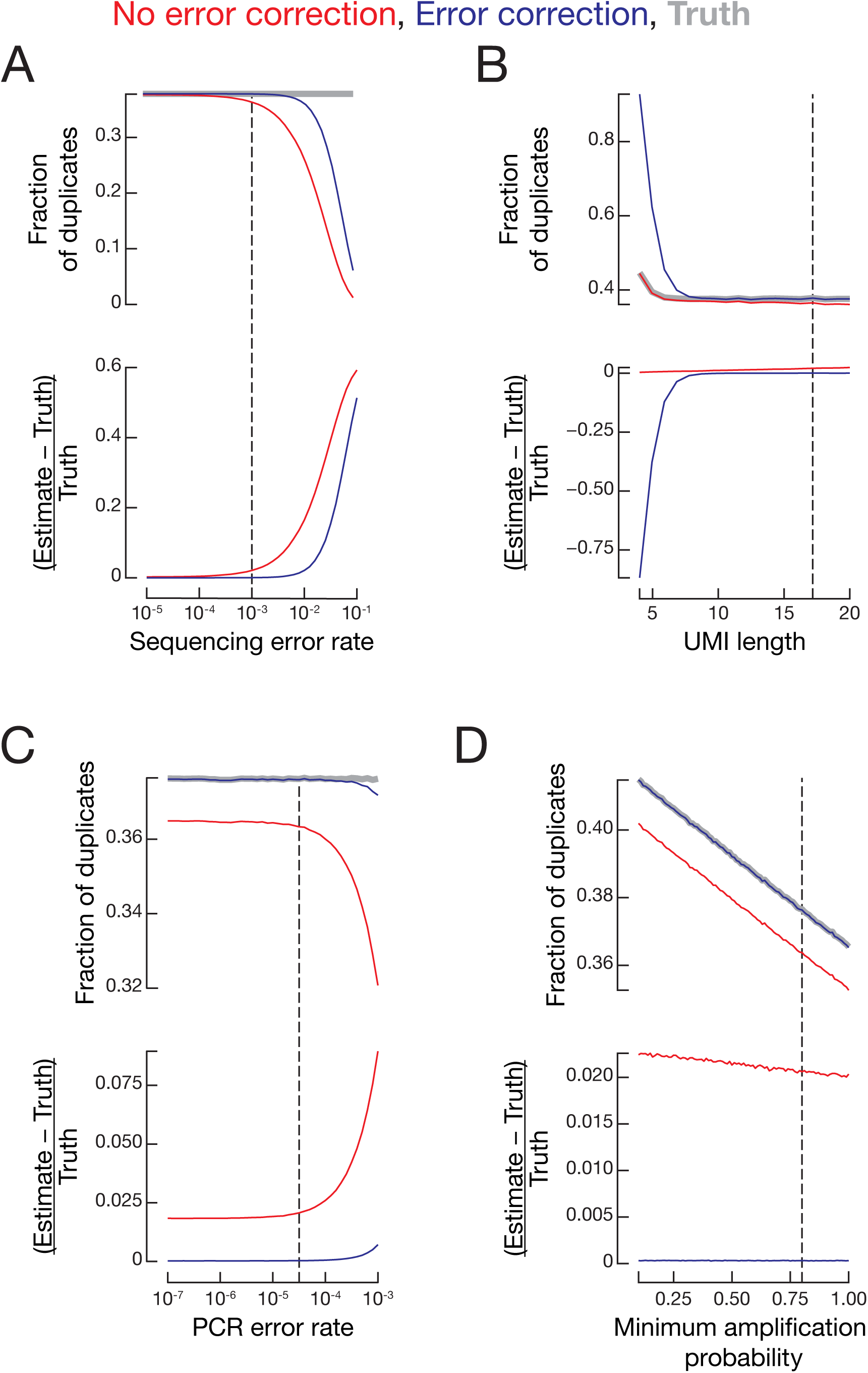
Accuracy and fraction of duplicates for simulated data varying (A) sequencing error rate, (B) UMI length, (C) PCR error rate, or (D) minimum amplification probability. Each dotted line indicates the value for this parameter used in other simulations.

Under some extreme conditions, correcting UMI errors yields substantially better results. For example, if we modify PCR error rate in the baseline condition from the default 3×10^-5^ (Lundberg et al., 1991; Zhou et al., 1991; Flaman et al., 1994) to 10^-3^, correcting UMI errors still yields a fraction of duplicates (37.2 ± 3.2%) very close to the truth (37.2 ± 3.1%), while not correcting the errors underestimates the fraction of duplicates (32.1 ± 3.5%). In conclusion, error-correction for UMIs consistently, albeit slightly, improves PCR duplicate identification. Therefore, we performed error correction for all following analyses.

### Removing PCR duplicates without using UMIs is fundamentally flawed

Does the common practice of removing PCR duplicates without UMIs improve the quantification of both long and short transcripts and in particular, of small RNAs such as microRNAs or piRNAs, which collectively originate from a small portion of the genome? We compared PCR duplicate identification using UMIs together with mapping coordinates of the reads to the conventional approach of using coordinates alone.

When only mapping coordinates were used (RNA-seq data from eight mouse tissues) (Supplemental Table S1A), 16.4%–44.5% RNA-seq reads were determined to be PCR duplicates, whereas using UMI information in conjunction with coordinates identified only 1.89%–10.67% as duplicates. That is, the majority of reads mapping to identical coordinates were in fact not PCR duplicates but rather from distinct starting molecules that should be counted for transcript abundance. The situation is even worse for small RNA-seq data, when only small RNA sequences were used, the majority (56.0%–76.8%) of reads were flagged as PCR duplicates and therefore excluded from analysis. In contrast, when UMI information was used together with the sequences of reads, just 1.05%–13.6% of reads were determined to be duplicates. Thus, most of the identical reads in RNA-seq and small RNA-seq are biologically real and not PCR duplicates, consistent with the view that small RNAs, which tend to come from precisely the same small genomic regions, can easily be mistaken for PCR duplicates when UMI information is not used. Moreover, the assumption that common mapping coordinates indicate PCR duplicates becomes increasingly problematic as sequencing depth increases, because the chance of observing two identical reads that legitimately derive from different molecules before PCR also increases.

We further tested whether PCR duplicate removal using only mapping coordinates is appropriate for transcript quantification (Fig. 5A). The conventional method underestimated the abundance of 119 transcripts by 1.25 fold or more: removing PCR duplicates based only on coordinates is too aggressive. These 119 transcripts are significantly shorter (median length = 602 nt) and more highly expressed (median abundance = 200 FPKM) than the other transcripts (median length = 1,620 nt; median abundance = 13.2 FPKM; Wilcoxon rank sum test *p* values = 2.22 × 10^-44^ and 1.80 × 10^-59^, respectively) (Fig. 5B). Thus, overestimation of PCR duplicates without UMIs reflects (1) a higher tendency of short transcripts to produce identical fragments due to more limited possibilities in fragmentation, and (2) a higher tendency of highly expressed genes to produce identical fragments. We conclude that removing PCR duplicates solely by mapping coordinates introduces substantial bias and that UMIs allow more accurate quantification of PCR duplicates and transcript abundance.

**Figure 5.**
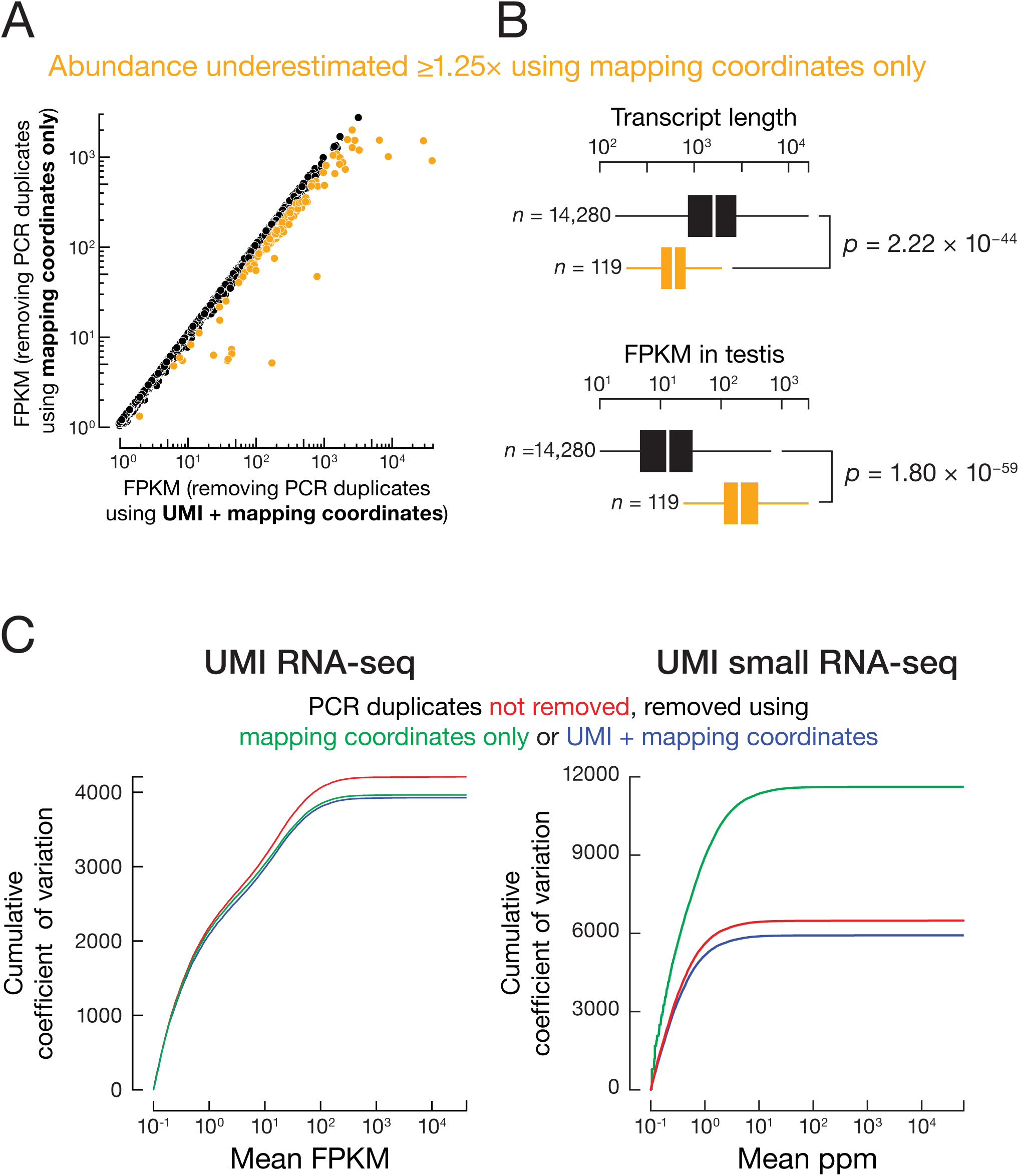
(A) Transcript abundance (FPKM) calculated by removing PCR duplicates using only mapping coordinates compared to using mapping coordinates and UMIs. (B) Using only mapping coordinates significantly biases against abundant and short genes. Outliers omitted. Wilcoxon rank sum test; n, number of genes in each group. (C) Relationship between cumulative coefficient of variation and transcript abundance.

### UMIs improve data reproducibility

One metric for evaluating the quality of experimental data is the reproducibility between technical replicates. We evaluated how UMIs affect the reproducibility of transcript quantification using five libraries generated using the same sample of total mouse testis RNA, but with gradually decreasing amounts of starting RNA and correspondingly increasing numbers of PCR cycles: 4 μg (8 PCR cycles), 2 μg (9 PCR cycles), 1 μg (10 PCR cycles), 500 ng (11 PCR cycles), 125 ng (13 PCR cycles) (Supplemental Table S1A). We then analyzed the data sets treating PCR duplicates using one of three approaches: (1) no PCR duplicates were removed; (2) PCR duplicates were removed using the conventional approach of identical genomic locations; and (3) PCR duplicates were removed using UMIs together with mapping coordinates. We compared the three approaches by calculating coefficients of variation (CV = S.D. / mean) for transcript abundance across the five RNA-seq libraries. Compared to removing no duplicates, removing duplicates according to their mapping coordinates decreased the total CV by 5.80% (from 4,210 to 3,960), while using UMIs with mapping coordinates decreased the total CV by 6.67% (from 4,210 to 3,930) (Fig. 5C). For example, when two RNA-seq libraries (125 ng with 12 PCR cycles and 1 μg with 10 PCR cycles) were compared, the number of transcripts whose abundance differed by ≥25% decreased when duplicates were removed (1,880 without duplicate removal, 1,503 removing duplicates by genomic coordinates, and 1,415 removing duplicates using UMIs). We conclude that removing PCR duplicates, using mapping coordinates alone or together with UMIs, improves the precision of transcript quantification.

Next, we evaluated the performance of these three approaches for a series of small RNA-seq libraries (starting material 39–5,000 ng). Compared to removing no duplicates, using UMIs to remove duplicates decreased the total CV by 8.72% (Fig. 5C). Surprisingly, removing duplicates according to their mapping coordinates alone increased CV by 79.1% (from 6,490 to 11,620) (Fig. 5C). For example, between two small RNA-seq libraries in this series, one generated from 150 ng and the other from 1 μg of the same total RNA sample, genomic loci (piRNA genes and GENCODE-annotated genes) whose small RNA abundance differed by ≥25% decreased 8.30% when duplicates were removed using UMIs (from 2,613 to 2,396 genes). In contrast, when duplicates were removed using solely mapping coordinates, the number of such irreproducible genes increased by 159% (6,762 genes). These results show that removing PCR duplicates with UMIs leads to more consistent quantification across libraries, whereas removing duplicates without UMIs is overly aggressive and decreases the reproducibility of small RNA-seq experiments.

### PCR cycles alone do not determine the frequency of PCR duplicates

It is widely accepted that the number of PCR cycles used to amplify the initial cDNA is the major cause of PCR duplicates in sequencing libraries (Andrews et al., 2016). We sought to test this assumption and to identify other experimental contributing factors. As described above, we performed computer simulations to test the impact of UMI error correction on PCR duplicate detection. We considered seven parameters that could impact the level of PCR duplicates during an RNA-seq or small RNA-seq experiment. Assuming that we have performed UMI error correction, we now examine in detail these seven parameters for their impact on the level of PCR duplicates.

Four of the parameters—PCR amplification efficiency, PCR error rate, sequencing error rate, and UMI length—are specified by the experimental reagents and sequencing platform and typically not adjusted from experiment to experiment. Our simulation results indicate that varying the sequencing error rate, the PCR error rate, or the UMI length around their default values in the baseline condition (i.e., within the ranges stipulated by experimental settings) did not have a significant effect on the faction of PCR duplicates (the blue line is flat around the dashed vertical line in Fig. S1A–C, top panels). In comparison, PCR efficiency had a measurable effect (the blue line in the top panel of Fig. S1D reveals a negative correlation with PCR efficiency). This is because that at lower PCR efficiency, some molecules are less likely to be amplified and become underrepresented, causing a decrease in library complexity and correspondingly higher fractions of PCR duplicates.

The other three parameters—the number of initial molecules, the number of molecules sequenced (i.e., sequencing depth), and the number of PCR cycles—are often adjusted to meet specific experimental conditions. Our simulations revealed that a change in PCR cycle number alone only minimally affected the fraction of PCR duplicates (the blue line in the top-left panel of Fig. 4 is nearly flat around the dashed vertical line), because the starting molecules of the original pool are proportionally propagated to the final library (Head et al., 2014). In contrast, decreasing the number of initial molecules or increasing the number of molecules sequenced sharply raised the frequency of PCR duplicates (Figure 4, two top-right panels).

We further tested these findings using experimental datasets. We first analyzed a set of five UMI RNA-seq libraries made with gradually decreasing amounts of starting RNA and correspondingly increasing numbers of PCR cycles: 4 μg (8 cycles), 2 μg (9 cycles), 1 μg (10 cycles), 500 ng (11 cycles), 125 ng (13 cycles) (Supplemental Table S1A). We observed that less starting RNA and correspondingly more PCR amplification resulted in higher fractions of PCR duplicates (Fig. 6A). For example, the 125 ng, 13-cycle library yielded 10.7% (median over 43,432 genes) PCR duplicates, while the 4 μg, 8-cycle library made by the same procedure contained only 1.79% PCR duplicates. Similarly, analysis of UMI small RNA-seq libraries generated from 39 ng (30 cycles) to 5 μg (16 cycles) total RNA (Supplemental Table S1A) revealed that starting with less RNA caused higher fractions of PCR duplicates (Fig. 6A).

**Figure 6.**
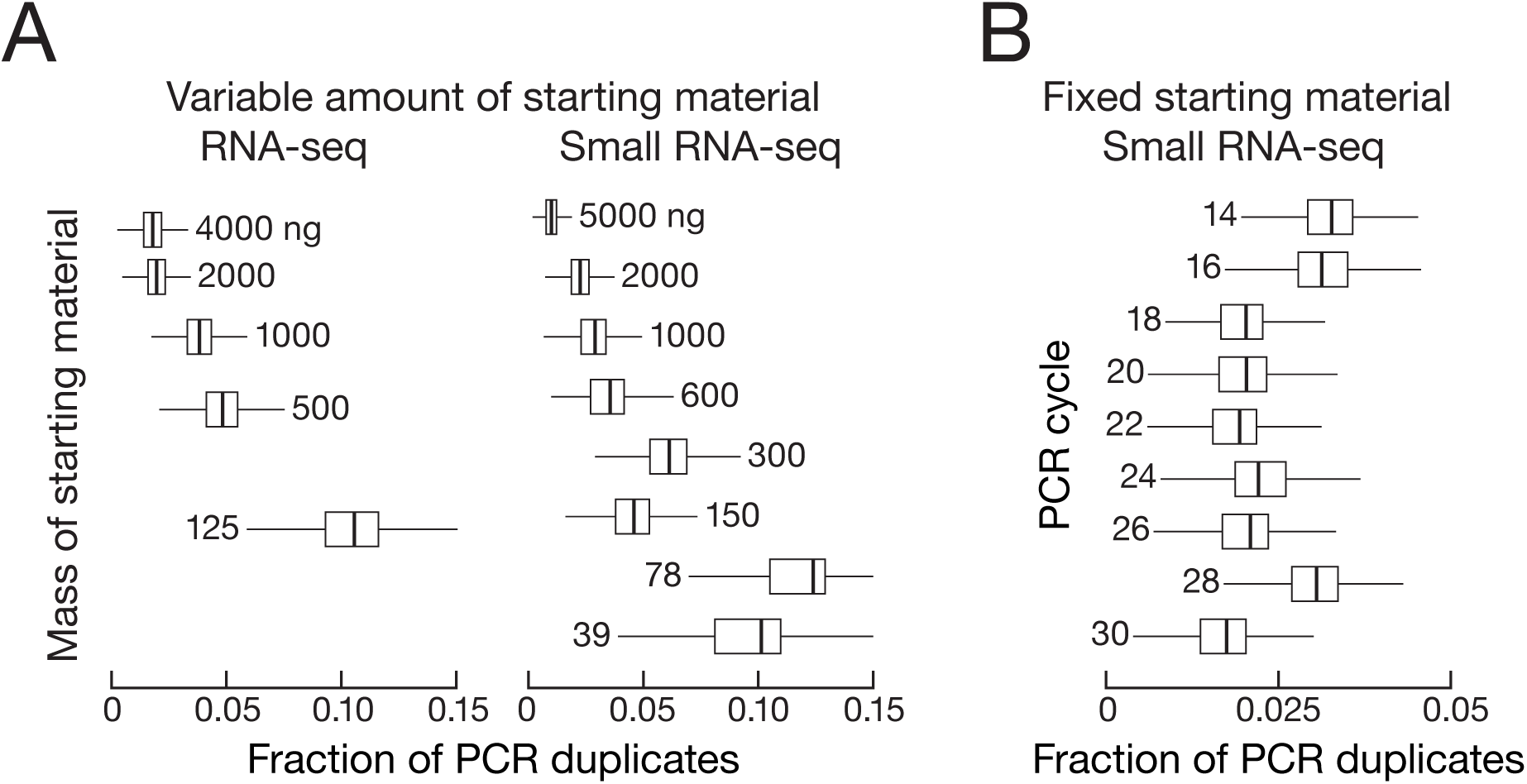
Fraction of PCR duplicates across genes for (A) a series of UMI RNA-seq and small RNA-seq libraries made with different amount of starting materials, and (B) a series of UMI small RNA-seq libraries all made with 5μg of total mouse testis RNA and with an increasing number of PCR cycles.

Simulations argue that the increase in PCR duplicates is not a consequence of greater PCR amplification but rather is caused by the use of lower starting material. To test this idea, we analyzed a second set of nine UMI small RNA-seq libraries, all generated from 5 μg total RNA from the same mouse testis, but amplified using 14 to 30 PCR cycles (Supplemental Table S1A). Consistent with the simulations, these libraries did not show a discernable trend between fraction of PCR duplicates and the number of PCR cycles (Fig. 6B). Thus, the higher fraction of PCR duplicates observed in libraries made from low amounts of RNA followed by high PCR cycle numbers more likely reflects the reduced complexity of the starting pool, rather than the increased number of PCR cycles. Together, our simulated and experimental data demonstrate that less starting RNA or higher sequencing depth, but not more PCR cycles *per se*, accounts for the frequency of PCR duplicates.

## Discussion

We have described experimental protocols and computational methods that, by incorporating UMIs into standard procedures, allow accurate PCR duplicate removal from RNA-seq and small RNA-seq data. Our approach increases reproducibility and decreases noise in sequencing libraries generated using a broad range of starting RNA amount and number of PCR cycles, enabling accurate quantification of the abundance of both long and short RNAs. We tested the importance of a key aspect of data processing—error correction for UMIs–and showed that under typical experimental conditions for bulk sequencing (represented by dotted lines in Fig. 4; Supplemental Fig. S1), correcting or not correcting errors in the UMI sequences has little absolute effect on PCR duplicate quantification. However, sequencing libraries made from a small number of cells, amount of tissue, or amount of RNA, have become increasingly common (Stegle et al., 2015), and they are more severely affected by PCR duplicates. Single-cell sequencing poses three specific challenges for PCR duplicate removal. First, it uses a limited amount of starting RNA, causing too low library complexity. Second, the ongoing discovery of new species of non-coding RNAs, many poorly understood, increases the number of species being measured, requiring longer UMIs. Finally, the increasingly high sequencing depth provided by advances in technology increases both the number of species that can be detected and the background noise. Together, these three factors make PCR duplicate measurement without UMI error correction especially problematic for single-cell sequencing. Our UMI approach should be directly applicable to single-cell RNA-seq. Error correction for UMIs mitigates these challenges by improving PCR duplicate identification.

The two most widely used computational tools for PCR duplicate removal, Picard MarkDuplicates (http://broadinstitute.github.io/picard/) and SAMtools rmdup (Li et al., 2009) rely only on the mapping coordinates of sequencing reads. Our data suggest that most identical reads reflect biological reality. Thus, removing PCR duplicate reads using only mapping coordinates erroneously eliminates many usable reads, particularly those produced from short transcripts and small RNAs.

The eight mouse tissues we analyzed span a range of transcriptome complexity: previous analyses showed that the mouse testis transcriptome contains ∼18,700 autosomal protein-coding transcripts, ∼8,600 non-coding RNAs, and ∼31.7 Mb of intergenic RNA, while the liver transcriptome contains only ∼15,500 autosomal protein-coding transcripts, ∼1,000 non-coding RNAs, and ∼7.2 Mb of intergenic RNA (Soumillon et al., 2013). Among the eight mouse tissues we tested, removing duplicate reads based on only mapping coordinates eliminates many biologically meaningful reads even when the libraries were made using ample starting RNA and optimal experimental conditions. Given the anti-correlation between RNA complexity and PCR duplicate occurrence, UMIs will improve the accuracy of comparing long or small RNA abundance across different tissues or cell types. Short RNAs, such as miRNAs and piRNAs, as well as highly abundant transcripts are particularly susceptible to underestimation by the conventional mapping coordinate method of PCR duplicate removal.

Our UMI approach builds on well-established protocols, requiring few changes in the procedures and little additional cost. We expect UMI analysis to be particularly useful when sequencing RNAs derived from a limited number of genomic loci, such as CaptureSeq (Mercer et al., 2014) and CAGE-seq (Carninci et al., 2006). Our approach can theoretically be adapted to any sequencing technique using synthetic oligonucleotide adapters. For example, sequencing immunoprecipitated chromatin (ChIP-seq) and the alternative CUT&RUN survey the genomic regions bound by proteins of interest (Park, 2009; Skene and Henikoff, 2017). The CUT&RUN method uses a nuclease to achieve more precise chromatin cleavage than the conventional ChIP-seq procedure, which utilizes sonication to randomly shear the DNA. Therefore, the likelihood of yielding identical reads also increases for CUT&RUN. By nature, protein-bound fragments also map to a smaller portion of genomic positions than RNA-seq reads. UMIs can improve discovery of protein binding sites by minimizing noise. Similarly, degradome sequencing profiles the 5′ ends of 3′ cleaved RNA products (Addo-Quaye et al., 2008); incorporating UMIs will enable precise quantification of cleaved RNA abundance.

## Methods

### Ribosomal RNA depletion for RNA-seq

Total RNA was extracted from tissues using the mirVana kit (ThermoFisher Scientific, Waltham, MA, USA) following manufacturer’s instructions. The ribosomal RNA depletion method was adapted from previously published protocols for human samples (Morlan et al., 2012; Adiconis et al., 2013). One hundred and eighty-six 50 nt-long DNA oligonucleotides complementary to the entire sequences of mouse 18S, 28S, 5S, and 5.8S rRNAs, and mitochondria 16S rRNA and 16S rRNA precursor were used at 0.5 μM (f.c.) for each oligonucleotide. Total mouse testis RNA was incubated with 1 μL of the DNA oligonucleotide mixture per 1 μg, and rRNA oligonucleotide hybridization buffer (100mM Tris-Cl pH 7.4, 200 mM NaCl) was added to make up to 10 μL. Oligonucleotide hybridization was carried out by heating the sample at 95°C for 3 min, then slowly cooling it down (−0.1°C/second) to 22°C in a thermocycler. The reaction was further incubated at 22°C for 5 min before being placed on ice. Thermostable RNase H (Lucigen, Middleton, MA) was added (5 U/μg total RNA), and the reaction adjusted to 50 mM Tris-Cl pH 7.4, 100 mM NaCl, and 20 mM MgCl_2_. and incubated at 45°C for 30 min. After DNase treatment with Turbo DNase (1 μL/μg total RNA) according to the manufacturer’s instructions, the rRNA-depleted RNA was purified using RNA Clean & Concentrator-5 (Zymo Research, Irvine, CA, USA).

### Sequencing

RNA-seq library generation was similar to previously published (Zhang et al., 2012), except for using UMI-containing adapters. Briefly, first strand cDNA was generated using ribosomal-depleted, fragmented total RNA. Resulting cDNA was incubated with a mixture of three sets of UMI-containing adapters, each carrying a distinct consensus sequence as described in the results for adapter ligation (Fig. 1B–C). The cDNA fragments ligated to adapters were amplified by PCR, and the length distributions and quality of the resulting libraries were analyzed by Agilent 2100 Bioanalyzer (Agilent Genomics, Santa Clara, MA, USA). The libraries were quantified using the KAPA library quantification kit (KAPA Biosystems, Wilmington, MA, USA) and sequenced using NextSeq 500 (Illumina, San Diego, CA, USA) paired-end sequencing.

Small RNA-seq library preparation was as previously described (Ghildiyal et al., 2008; Seitz et al., 2008). Briefly, 18–35 nt small RNAs were size-selected by polyacrylamide gel electrophoresis using RNA markers. Small RNAs were first ligated to 3′ DNA adapters with adenylated 5′ and dideoxycytosine-blocked 3′ ends. These contained UMIs in 3 nt-blocks of random nucleotides separated by pre-defined 3 nt consensus sequences (NNN-GTC-NNN-TAG-NNN, Fig. 2B). The ligated products were purified by polyacrylamide gel electrophoresis and ligated to a mixed pool of equimolar amount of 5′ RNA adapters containing UMIs in 3 nt-blocks of random nucleotides and one of the two distinct consensus sequence sets (NNN-CGA-NNN-UAC-NNN and NNN-AUC-NNN-AGU-NNN). The length distributions and quality of the resulting libraries were analyzed by Agilent 2100 Bioanalyzer. The libraries were quantified using the KAPA library quantification kit and sequenced using NextSeq 500 single-end sequencing.

### Bioinformatics

Simulation procedure was performed according to (Smith et al., 2017). Briefly, we simulated 7 parameters: PCR amplification probability, PCR and sequencing error rates, UMI length, number of initial molecules, number of sequenced molecules, and PCR cycle numbers, by varying one parameter and keeping other parameters constant. For each combination of the 7 parameters, we performed 10,000 replicates. UMI error correction was implemented as described in (Smith et al., 2017), except that we used read sequences instead of genomic coordinates when determining PCR duplicates for small RNA-seq. We used NetworkX (Hagberg et al., 2008) for graph-related algorithms, and pysam (https://github.com/pysam-developers/pysam) for handling SAM/BAM files. Reads were mapped to the mouse mm10 genome as described in (Han et al., 2015). When reads were analyzed without UMIs, PCR duplicates were identified using Picard (https://github.com/broadinstitute/picard).

### Data access

The tools developed for handling UMIs in our RNA-seq and small RNA-seq data can be found at https://github.com/weng-lab/umitools, and via PyPI (package: umitools). RNA-seq and small RNA-seq data have been deposited in the NCBI SRA under the accession number PRJNA416930.

## Acknowledgements

We thank members of the Weng and Zamore laboratories for helpful discussions and comments on the manuscript; This work was supported in part by National Institutes of Health grants P01HD078253 and R37GM062862 to PDZ and HD078253 to ZW.

## Author contributions

YF, Bioinformatics analysis, Writing; PHW, Protocol design, RNA-seq, small RNA-seq, Writing; ZW, Project design and management, Writing; PDZ, Project conception, design and management, Writing.

---

**Supplemental Table S1.** Mapping and UMI statistics of (A) RNA-seq and (B) small RNA-seq data generated in this study.

## References

Addo-Quaye C, Eshoo TW, Bartel DP, Axtell MJ. 2008. Endogenous siRNA and miRNA targets identified by sequencing of the Arabidopsis degradome. Curr Biol 18: 758–762.

Adiconis X, Borges-Rivera D, Satija R, DeLuca DS, Busby MA, Berlin AM, Sivachenko A, Thompson DA, Wysoker A, Fennell T, Gnirke A, Pochet N, Regev A, Levin JZ. 2013. Comparative analysis of RNA sequencing methods for degraded or low-input samples. Nat Methods 10: 623–629.

Andrews KR, Good JM, Miller MR, Luikart… G. 2016. Harnessing the power of RADseq for ecological and evolutionary genomics. Nature Reviews …

Aravin A, Gaidatzis D, Pfeffer S, Lagos-Quintana M, Landgraf P, Iovino N, Morris P, Brownstein MJ, Kuramochi-Miyagawa S, Nakano T, Chien M, Russo JJ, Ju J, Sheridan R, Sander C, Zavolan M, Tuschl T. 2006. A novel class of small RNAs bind to MILI protein in mouse testes. Nature 442: 203–207.

Aravin AA, Sachidanandam R, Girard A, Fejes-Toth K, Hannon GJ. 2007. Developmentally regulated piRNA clusters implicate MILI in transposon control. Science 316: 744–747

Bose S, Wan Z, Carr A, Rizvi AH, Vieira G, Pe’er D, Sims PA. 2015. Scalable microfluidics for single-cell RNA printing and sequencing. Genome Biol 16: 120.

Brennecke J, Aravin AA, Stark A, Dus M, Kellis M, Sachidanandam R, Hannon GJ. 2007. Discrete small RNA-generating loci as master regulators of transposon activity in Drosophila. Cell 128: 1089–1103.

Carninci P, Sandelin A, Lenhard B, Katayama S, Shimokawa K, Ponjavic J, Semple CA, Taylor MS, Engström PG, Frith MC, Forrest AR, Alkema WB, Tan SL, Plessy C, Kodzius R, Ravasi T, Kasukawa T, Fukuda S, Kanamori-Katayama M, Kitazume Y, Kawaji H, Kai C, Nakamura M, Konno H, Nakano K, Mottagui-Tabar S, Arner P, Chesi A, Gustincich S, Persichetti F, Suzuki H, Grimmond SM, Wells CA, Orlando V, Wahlestedt C, Liu ET, Harbers M, Kawai J, Bajic VB, Hume DA, Hayashizaki Y. 2006. Genome-wide analysis of mammalian promoter architecture and evolution. Nat Genet 38: 626–635.

Cha RS, Thilly WG. 1993. Specificity, efficiency, and fidelity of PCR. PCR Methods Appl 3: S18–29.

Collins JE, Wali N, Sealy IM, Morris JA, White RJ, Leonard SR, Jackson DK, Jones MC, Smerdon NC, Zamora J, Dooley CM, Carruthers SN, Barrett JC, Stemple DL, Busch-Nentwich EM. 2015. High-throughput and quantitative genome-wide messenger RNA sequencing for molecular phenotyping. BMC Genomics 16: 578.

Dohm JC, Lottaz C, Borodina T, Himmelbauer H. 2008. Substantial biases in ultra-short read data sets from high-throughput DNA sequencing. Nucleic Acids Res 36: e105.

Flaman JM, Frebourg T, Moreau V, Charbonnier F, Martin C, Ishioka C, Friend SH, Iggo R. 1994. A rapid PCR fidelity assay. Nucleic Acids Res 22: 3259–3260.

Fu GK, Hu J, Wang PH, Fodor SP. 2011. Counting individual DNA molecules by the stochastic attachment of diverse labels. Proc Natl Acad Sci U S A 108: 9026–9031.

Fu GK, Wilhelmy J, Stern D, Fan HC, Fodor SP. 2014a. Digital encoding of cellular mRNAs enabling precise and absolute gene expression measurement by single-molecule counting. Anal Chem 86: 2867–2870.

Fu GK, Xu W, Wilhelmy J, Mindrinos MN, Davis RW, Xiao W, Fodor SP. 2014b. Molecular indexing enables quantitative targeted RNA sequencing and reveals poor efficiencies in standard library preparations. Proc Natl Acad Sci U S A 111: 1891–1896.

Ghildiyal M, Seitz H, Horwich MD, Li C, Du T, Lee S, Xu J, Kittler EL, Zapp ML, Weng Z, Zamore PD. 2008. Endogenous siRNAs derived from transposons and mRNAs in Drosophila somatic cells. Science 320: 1077–1081.

Girard A, Sachidanandam R, Hannon GJ, Carmell MA. 2006. A germline-specific class of small RNAs binds mammalian Piwi proteins. Nature

Hagberg A, Swart P, Chult DS. 2008. Exploring network structure, dynamics, and function using NetworkX. permalink.lanl.gov

Han BW, Wang W, Zamore PD, Weng Z. 2015. piPipes: a set of pipelines for piRNA and transposon analysis via small RNA-seq, RNA-seq, degradome‐ and CAGE-seq, ChIP-seq and genomic DNA sequencing. Bioinformatics 31: 593–595.

Hayashi R, Handler D, Ish-Horowicz D, Brennecke J. 2014. The exon junction complex is required for definition and excision of neighboring introns in Drosophila. Genes Dev 28: 1772–1785.

Head SR, Komori HK, LaMere SA, Whisenant T, Van Nieuwerburgh F, Salomon DR, Ordoukhanian P. 2014. Library construction for next-generation sequencing: overviews and challenges. Biotechniques 56: 61–64.

Houwing S, Kamminga LM, Berezikov E, Cronembold D, Girard A, van den Elst H, Filippov DV, Blaser H, Raz E, Moens CB, Plasterk RH, Hannon GJ, Draper BW, Ketting RF. 2007. A role for Piwi and piRNAs in germ cell maintenance and transposon silencing in Zebrafish. Cell 129: 69–82.

Illumina. 2014. Illumina: Low-Diversity Sequencing on the Illumina HiSeq^®^ Platform. Technical Note: DNA Sequencing

Illumina. 2016. Illumina: NextSeq^®^ 500 System Guide. Technical Note: DNA Sequencing

Islam S, Zeisel A, Joost S, La Manno G, Zajac P, Kasper M, Lönnerberg P, Linnarsson S. 2014. Quantitative single-cell RNA-seq with unique molecular identifiers. Nat Methods 11: 163–166.

Kivioja T, Vähärautio A, Karlsson K, Bonke M, Enge M, Linnarsson S, Taipale J. 2011. Counting absolute numbers of molecules using unique molecular identifiers. Nat Methods 9: 72–74.

Lau NC, Lim LP, Weinstein EG, Bartel DP. 2001. An abundant class of tiny RNAs with probable regulatory roles in Caenorhabditis elegans. Science

Li H, Handsaker B, Wysoker A, Fennell T, Ruan J, Homer N, Marth G, Abecasis G, Durbin R, 1000 GPDPS. 2009. The Sequence Alignment/Map format and SAMtools. Bioinformatics 25: 2078–2079.

Li XZ, Roy CK, Dong X, Bolcun-Filas E, Wang J, Han BW, Xu J, Moore MJ, Schimenti JC, Weng Z, Zamore PD. 2013. An ancient transcription factor initiates the burst of piRNA production during early meiosis in mouse testes. Mol Cell 50: 67–81.

Lundberg KS, Shoemaker DD, Adams MW, Short JM, Sorge JA, Mathur EJ. 1991. High-fidelity amplification using a thermostable DNA polymerase isolated from Pyrococcus furiosus. Gene 108: 1–6.

Macosko EZ, Basu A, Satija R, Nemesh J, Shekhar K, Goldman M, Tirosh I, Bialas AR, Kamitaki N, Martersteck EM, Trombetta JJ, Weitz DA, Sanes JR, Shalek AK, Regev A, McCarroll SA. 2015. Highly Parallel Genome-wide Expression Profiling of Individual Cells Using Nanoliter Droplets. Cell 161: 1202–1214.

Mercer TR, Clark MB, Crawford J, Brunck ME, Gerhardt DJ, Taft RJ, Nielsen LK, Dinger ME, Mattick JS. 2014. Targeted sequencing for gene discovery and quantification using RNA CaptureSeq. Nat Protoc 9: 989–1009.

Mitra A, Skrzypczak M, Ginalski K, Rowicka M. 2015. Strategies for achieving high sequencing accuracy for low diversity samples and avoiding sample bleeding using illumina platform. PLoS One 10: e0120520.

Mohn F, Sienski G, Handler D, Brennecke J. 2014. The rhino-deadlock-cutoff complex licenses noncanonical transcription of dual-strand piRNA clusters in Drosophila. Cell

Morlan JD, Qu K, Sinicropi DV. 2012. Selective depletion of rRNA enables whole transcriptome profiling of archival fixed tissue. PLoS One 7: e42882.

Park PJ. 2009. ChIP-seq: advantages and challenges of a maturing technology. Nat Rev Genet 10: 669–680.

Schirmer M, D’Amore R, Ijaz UZ, Hall N, Quince C. 2016. Illumina error profiles: resolving fine-scale variation in metagenomic sequencing data. BMC Bioinformatics 17: 125.

Seitz H, Ghildiyal M, Zamore PD. 2008. Argonaute loading improves the 5’ precision of both MicroRNAs and their miRNA* strands in flies. Curr Biol 18: 147–151.

Shiroguchi K, Jia TZ, Sims PA, Xie XS. 2012. Digital RNA sequencing minimizes sequence-dependent bias and amplification noise with optimized single-molecule barcodes. Proc Natl Acad Sci U S A 109: 1347–1352.

Skene PJ, Henikoff S. 2017. An efficient targeted nuclease strategy for high-resolution mapping of DNA binding sites. eLife 6: e21856.

Smith T, Heger A, Sudbery I. 2017. UMI-tools: modeling sequencing errors in Unique Molecular Identifiers to improve quantification accuracy. Genome Res 27: 491–499.

Soumillon M, Necsulea A, Weier M, Brawand D, Zhang X, Gu H, Barthès P, Kokkinaki M, Nef S, Gnirke A, Dym M, de Massy B, Mikkelsen TS, Kaessmann H. 2013. Cellular source and mechanisms of high transcriptome complexity in the mammalian testis. Cell Rep 3: 2179–2190.

Stegle O, Teichmann SA, Marioni JC. 2015. Computational and analytical challenges in single-cell transcriptomics. Nature Reviews Genetics 16: 133-45.

Yaari G, Kleinstein SH. 2015. Practical guidelines for B-cell receptor repertoire sequencing analysis. Genome Med 7: 121.

Yue F, Cheng Y, Breschi A, Vierstra J, Wu W, Ryba T, Sandstrom R, Ma Z, Davis C, Pope BD, Shen Y, Pervouchine DD, Djebali S, Thurman RE, Kaul R, Rynes E, Kirilusha A, Marinov GK, Williams BA, Trout D, Amrhein H, Fisher-Aylor K, Antoshechkin I, DeSalvo G, See LH, Fastuca M, Drenkow J, Zaleski C, Dobin A, Prieto P, Lagarde J, Bussotti G, Tanzer A, Denas O, Li K, Bender MA, Zhang M, Byron R, Groudine MT, McCleary D, Pham L, Ye Z, Kuan S, Edsall L, Wu YC, Rasmussen MD, Bansal MS, Kellis M, Keller CA, Morrissey CS, Mishra T, Jain D, Dogan N, Harris RS, Cayting P, Kawli T, Boyle AP, Euskirchen G, Kundaje A, Lin S, Lin Y, Jansen C, Malladi VS, Cline MS, Erickson DT, Kirkup VM, Learned K, Sloan CA, Rosenbloom KR, Lacerda de Sousa B, Beal K, Pignatelli M, Flicek P, Lian J, Kahveci T, Lee D, Kent WJ, Ramalho Santos M, Herrero J, Notredame C, Johnson A, Vong S, Lee K, Bates D, Neri F, Diegel M, Canfield T, Sabo PJ, Wilken MS, Reh TA, Giste E, Shafer A, Kutyavin T, Haugen E, Dunn D, Reynolds AP, Neph S, Humbert R, Hansen RS, De Bruijn M, Selleri L, Rudensky A, Josefowicz S, Samstein R, Eichler EE, Orkin SH, Levasseur D, Papayannopoulou T, Chang KH, Skoultchi A, Gosh S, Disteche C, Treuting P, Wang Y, Weiss MJ, Blobel GA, Cao X, Zhong S, Wang T, Good PJ, Lowdon RF, Adams LB, Zhou XQ, Pazin MJ, Feingold EA, Wold B, Taylor J, Mortazavi A, Weissman SM, Stamatoyannopoulos JA, Snyder MP, Guigo R, Gingeras TR, Gilbert DM, Hardison RC, Beer MA, Ren B, Mouse ENCODEC. 2014. A comparative encyclopedia of DNA elements in the mouse genome. Nature 515: 355–364.

Zhang Z, Theurkauf WE, Weng Z, Zamore PD. 2012. Strand-specific libraries for high throughput RNA sequencing (RNA-Seq) prepared without poly(A) selection. Silence 3: 9.

Zhang Z, Wang J, Schultz N, Zhang F, Parhad SS, Tu S, Vreven T, Zamore PD, Weng Z, Theurkauf WE. 2014. The HP1 homolog rhino anchors a nuclear complex that suppresses piRNA precursor splicing. Cell 157: 1353–1363.

Zhou YH, Zhang XP, Ebright RH. 1991. Random mutagenesis of gene-sized DNA molecules by use of PCR with Taq DNA polymerase. Nucleic Acids Research 19: 6052.

